# Facing the green threat: A waterflea’s defenses against a carnivorous plant

**DOI:** 10.1101/2021.10.19.464940

**Authors:** Sebastian Kruppert, Martin Horstmann, Linda C. Weiss, Elena Barmaeva, Nadja Kubitza, Simon Poppinga, Anna S. Westermeier, Thomas Speck, Ralph Tollrian

## Abstract

Water fleas of the family Daphniidae are keystone species in many lentic ecosystems and, as most abundant filter feeders, link the primary production to higher trophic levels. As a response to the high predatory pressures, water fleas have evolved a range of defenses, including inducible defenses against animal predators. Here we show in *Ceriodaphnia dubia* a first example of such defenses induced by the presence of a coexisting plant predator, i.e. the carnivorous southern bladderwort (*Utricularia australis*, Lentibulariaceae), which possesses ultrafast underwater suction traps. When the bladderwort is present, *C. dubia* shows changes in morphology, life-history and behavior. While the morphological and behavioral adaptations improve *C. dubia’s* survival rate in the presence of this predator, the life-history parameters likely reflect trade-offs for the defense. Our study demonstrates plant-induced animal defenses, implying their potential relevance in freshwater ecosystems and contributing to an overall yet underestimated biodiversity of inducible defenses.

**Open Research Statement:** Data is not finally prepared for upload yet. Once most fitting file types are determined and metadata is created we aim to upload all raw data as supporting information.

## Background

Members of the crustacean family Daphniidae represent some of the most abundant zooplankters in lentic freshwater ecosystems (1). As consumers of phytoplankton, they link primary production to higher trophic levels by falling prey to a variety of predators like other crustaceans, fish or insects (2,3). This seasonally high but variable predation risk favored the evolution of inducible defenses in some species of the Daphniidae. Inducible defenses are a form of phenotypic plasticity that decreases an organism’s vulnerability to specific predators (for reviews see Tollrian & Harvell 1999; Weiss *et al*. 2012; Weiss & Tollrian 2018; Riessen & Gilbert 2019). These defenses range from alterations in morphology, or life history parameters to behavior. Many defenses are predator-specific and adapted to counter the respective predator. For example, a vast diversity of striking morphological defenses has been described. They include rather minute structures like ‘neckteeth’ expressed by *Daphnia pulex* (8–10) and the medium sized ‘crown of thorns’ in *D. atkinsoni* (11) or large morphological changes like the helmets of *D. cucullata* and *D. lumholtzi* (12,13) and the crests of *D. longicephala* (14–16). Further, alterations in the carapace architecture and its mechanical properties have been reported (17–19). In the presence of visual hunters like fish, some *Daphnia* species alter their life history and shift resources from somatic growth to reproduction (20,21). Species of *Daphnia* predated by invertebrates display the opposite strategy by accelerating their somatic growth. This way prey overcomes the predator’s gape limit, at the expense of population growth rate (22–25). A well-studied behavioral defense strategy is the diel vertical migration of zooplankton. Here, *Daphnia* avoid visual predators by residing in deeper waters during the day and ascent into shallow nutrient-rich strata for grazing during the night when these predators are hampered by low light levels (26). Also, changes in swimming behavior have been reported, like predator-induced increase or decrease in individual swimming speed (for a review see Langer *et al*. 2019).

Existing literature on inducible defenses in Daphniidae focusses on responses against animal predators overlooking carnivorous plants. One exception is a study by Havel and Dodson (28) that included *Utricularia* spec. but observed no induction of morphological defenses in *D. retrocurva*.

The southern bladderwort (*Utricularia australis*), native to Central Europe, is a naturally coexisting predator of many different *Daphnia* species including *Ceriodaphnia dubia* (29–31). With its ultrafast suction traps, it can catch prey within ∼5 ms, leaving little escape chances (32). Water is actively pumped out of the trap lumen via specialized glands (33), creating sub-ambient pressure inside the trap (34–36). If triggered, *C. dubia* is sucked into the trap with a speed of up to 4 m/s (32). The trap resets in about 15-30 min after suction and continues to catch further prey until the trap is full. With the help of these highly efficient traps the plant acquires a substantial nutrient supply (32,36–38). According to their spatial dimensions, many species within the Daphniidae family fit into the suction traps and are therefore potential prey (39). In combination with seasonally high abundances of *U. australis*, and due to the fact that each plant can possess several hundreds to thousands of traps, it may pose a severe threat on daphniid populations (31). In fact, trap content analysis prove Daphniidae to be a substantial portion of the southern bladderwort’s prey spectrum (32). In this context, we hypothesized that *Daphnia* may have evolved mechanisms reducing this predation pressure.

Using high-resolution 3D morphometrics (40), we investigated morphological changes in *C. dubia* as adaptive responses to the presence of *U. australis*. Additionally, we analyzed life history shifts and behavioral alterations as a possible response of *C. dubia* to the plant’s presence. Furthermore, we analyzed the bladderwort’s capture efficiency for control and exposed (defense-induced) *C. dubia* in order to determine the protective effect of the displayed defensive strategies.

## Materials and Methods

### Study Design

In order to depict a naturally occurring predator-prey system we started this study by identifying local ponds containing *U. australis* alongside several Daphniidae species in the field. We subsequently performed a trap analysis to validate the co-occurring Daphniidae species as prey items of *U. australis*. From the resulting prey spectrum analysis (32) we chose *C. dubia* as our candidate for the present study due to its high abundance in the pond as well as in the *U. australis* traps. For validating our initial hypothesis that *C. dubia* has evolved inducible defenses against the coexisting bladderwort (*U. australis*), we adjusted controlled laboratory experiments initially developed for animal predators (e.g. Tollrian 1990, 1994; Weiss *et al*. 2016; Kruppert *et al*. 2017; Poppinga *et al*. 2017). Based on our experience with Daphniidae and their inducible defenses, we aimed for a sample size of 10 specimens for each experiment, as we expected any alterations to be detectable with this sample size (e.g. Tollrian 1990; Kruppert *et al*. 2016, 2017; Weiss *et al*. 2016; Horstmann *et al*. 2018). The first experiment was designed to verify whether *C. dubia* reacts on the presence of *U. australis* with alterations in morphology and life history. Using light microscopy, we measured morphometric (body length and body width) as well as life history parameters (number of egg-carrying females, clutch size) of initially juvenile *C. dubia* specimen in four different treatments (tap water control, non-threatening plant, fed *Utricularia*, unfed *Utricularia*) over a duration of 6 days. Based on the initial findings, we conducted follow-up experiments in order to identify behavioral alterations as well as to validate the alterations as being an effective defense to the bladderwort traps. All experiments are described in detail below. We did not exclude any data from the analysis and outliers where not predefined or treated differently in the analysis. Randomization, where conducted, was used to prevent influence of external factors (i.e. illumination) and no specific method for randomization was applied. Our study does not include any mode of blinding.

### Sampling of *C. dubia*

*C. dubia* individuals coexisting with *U. australis* in a fish-free, artificial pond complex in Gelsenkirchen, Germany (51°30’17.9”N 7°04’58.7”E) were sampled October 1^st^, 2015 and brought immediately to the lab for culturing.

### Cultures

#### Prey crustaceans (*Ceriodaphnia dubia*)

From the Gelsenkirchen pond samples, a clonal line of *C. dubia* (S04) was reared from a single female. This female and the subsequent offspring were cultured in 1 L beakers (J. Weck GmbH & Co. KG, Wehr-Öflingen, Germany) containing charcoal-filtered tap-water. A maximum of 100 animals was kept in the beakers by transferring supernumerary adults and neonates into new beakers. The beakers were regularly cleared of detritus, half of the water was exchanged monthly and *Acutodesmus obliquus* was added as food source *ad libitum*. The cultures were kept under stable conditions at 20°C +/- 1°C and a 16 h:8 h light to dark cycle.

#### Predator (the carnivorous plant *Utricularia australis*)

We used *U. australis* cultivated and used in prior experiments at the Botanical Garden of the University of Freiburg. The plants were cultivated in the Department of Animal Ecology, Evolution and Biodiversity of the Ruhr-University Bochum, Germany. Plants were kept in 50 L plastic aquaria filled with charcoal-filtered tap-water and positioned 60 cm beneath a light source consisting of four fluorescent tube lamps with 36 W each (Radium NL 36 W/840 Spectralux Plus cool white). The *U. australis* culture was kept under the same stable conditions as the *C. dubia* culture (at 20 °C +/- 1 °C and a 16 h:8 h light to dark cycle), and the plants were constantly growing and continuously producing new traps.

#### Trap entrance dimensions

To measure the predator’s gape size, twenty *U. australis* traps were dissected from the plant and imaged using a stereomicroscope (Olympus SZX16) with a digital camera (ColorView III digital imaging system) attached. The widths and heights of the trap entrances were measured via imaging software (Cell^D; Soft Imaging Solutions, SIS Olympus, Münster, Germany). As trap entrance width, we defined the shortest distance between opposite trap entrance walls, parallel to the threshold of the trap entrance margin (42). The height of the trap entrance is the line connecting threshold and trap door insertion and is therefore orthogonal to the width.

### Defense induction

In order to investigate *U. australis-*induced morphological and life-history defenses in *C. dubia*, we analyzed individuals from the earliest juvenile stages. To do so, we started the experiments with egg-carrying individuals in the last embryonic stage and measured the offspring individually every 24 hours throughout the following 6 days. We chose this ontogenetic stage because *Daphnia* is sensitive to predatory cues from the fourth embryonic stage onward (10). We conducted the experiment in a full factorial design consisting of four different treatments (n=10 each). We used two different treatments in order to control for the absence of plants (‘tap water control’) as well as for the presence of non-threatening plants by exposing *C. dubia* to an equal amount of coontails (*Ceratophyllum demersum*) as we used *U. australis* in the experimental treatments (see below) (‘*Ceratophyllum* control’). Coontails naturally occur together with *U. australis* (43) and *C. dubia*. As experimental treatments we conducted two induction setups where *C. dubia* was confronted with *U. australis*. In order to identify whether the biological activity is solely plant borne, we reared *C. dubia* together with bladderworts as one experimental treatment (‘unfed *Utricularia*’). In addition, we performed an experimental treatment in which *C. dubia* was exposed to bladderwort that were fed daily with 25 juvenile *C. dubia* (‘fed *Utricularia*’), as inducing agents are often associated with active feeding processes (44,45). All treatments were conducted in 1 L beakers (J. Weck GmbH & Co. KG, Wehr-Öflingen, Germany). To avoid direct predator contact and prevent the consumption of the test specimens in both predator treatments, we separated prey (*C. dubia*) and predator (*U. australis*) using net cages equipped with fine mesh widths of 125 µm (Hydrobios, Germany). Within the net cages, we placed the egg carrying *C. dubia* females. Plants (one shoot of 10-15 cm each) were placed outside the net cages and, depending on the treatment, were fed daily with 25 juvenile *C. dubia* (‘fed *Utricularia*’) or left unfed.

### Analysis of morphology and life history alterations

Once *C. dubia* females released their brood (i.e. approximately within 24 h), we removed the mothers leaving only the offspring in the net cages. Another 24 h later, we started to image the animals in a daily rhythm for 6 days in total by using a stereomicroscope (Olympus SZX16) equipped with a digital camera (ColorView III digital imaging system) and imaging software (Cell^D; Soft Imaging Solutions, SIS Olympus, Münster, Germany). We measured the following parameters in order to identify morphological and life-history alterations (see S-Table 1, 2 and 5): body length, body width, the number of egg carrying females and the average number of eggs deposited in the brood pouch. The body length was measured from the top of the compound eye to the point where the carapace converges into the tail-spine. Body width was measured at the broadest distance between ventral and dorsal perpendicular to the body length. In order to analyze the body width allometrically, we normalized it to the body length (normalized body width = body width/body length).

### 3D analysis of morphological alterations

In order to identify morphological alterations comprehensively, we conducted a three-dimensional analysis of control and plant-exposed *C. dubia* as described by Horstmann et al. (40). For that, we used *C. dubia* (n(control)=13; n(induced)=8) individuals from the ‘fed *Utricularia’* treatment on day 5 of the experimental period. The animals were stained using Congo red, scanned on a confocal laser scanning microscope and subsequently digitized as a surface image. These surface images were analyzed using a landmark based method (≈45.000 semi-landmarks per animal) and compared using a Procrustes-based analysis. For details please see Horstmann et al. 2018 (40).

### Analysis of behavioral defenses

#### Predator avoidance

We conducted a subsequent experiment that aimed to identify behavioral changes in *C. dubia* as a response to the presence of *U. australis*. We designed this experiment in order to test whether *C. dubia* avoids areas that are shadowed by plants based in dependence of their stage of alertness (either naïve or alerted by prior predator exposure). We used five different setups resulting from the combination of two different control treatments (‘control’ and ‘fed *Utricularia*’) and three different experimental scenarios (‘no plant’, ‘*Elodea*’ and ‘*Utricularia*’). The combination ‘fed *Utricularia*’ / ‘no plant’ was not included in our experiments. As specimens, we used five-day-old *C. dubia* that were either reared in ‘control’ or in ‘fed *Utricularia*’ beakers. The ‘no plant’ scenario with ‘control’ animals was used as behavioral baseline (‘tap water control’). Avoidance behavior of the two treatments was tested in the two environmental scenarios; a control condition with the non-carnivorous plant *Elodea canadensis* and a test treatment with the carnivorous plant *U. australis*. We wanted to test for external factors affecting behavior (e.g. inhomogeneous light conditions, as a result of plant associated shading) and used exposure to *Elodea canadensis* as a comparison to the control condition without any plants because *E. canadensis* shows strong similarity to *U. australis* in terms of shadowing. Their color and whorl morphology give them an *Utricularia-*like appearance. The plant treatments were conducted using a single shoot of *U. australis* or *E. canadensis* respectively. For each treatment, we placed 20 five-day-old *C. dubia* in 2 L plastic tanks (ca. 18 cm × 13 cm × 11.5 cm, Savic, Kortrijk, Belgium) filled with charcoal-filtered tap-water and according to the experimental condition, *U. australis* or *E. canadensis* randomly positioned on either side of the tank. The plants were kept on one side of the tank with a spacer positioned centrally in the respective tank, fixing the floating plants. All five different experimental setups (‘tap water control’, ‘control vs. *Elodea*’, ‘fed *Utricularia* vs. *Elodea*’, ‘control vs. *Utricularia*’, ‘fed *Utricularia* vs. *Utricularia*’) were started simultaneously and monitored in parallel. The experiment was repeated 10 times. For documentation of the animals’ positions, the tanks were divided into 18 equally sized sections (each approx. 3×4.3 cm) by superimposing a grid with three rows and six columns on the tanks’ fronts. Three of these columns did not contain plants, three columns contained plants. For homogenous light conditions and to avoid light reflections, we installed a single fluorescent tube lamp above each tank, (fluorescent tube lamp, Radium NL 36 W/840 Spectralux Plus cool white). This setup provided uniform light over the whole surface and prevented shadows. Furthermore, the treatments were randomly permutated between the tanks in order to exclude position-dependent effects (e.g. whether there were neighbouring tanks or not). We started the experiment by introducing the 20 five-day-old *C. dubia* after acclimation for 30 min to the new environment as used in comparable studies (27,46). We manually documented the distribution pattern of *C. dubia* in the sections of the tank every 15 minutes for a total duration of 60 minutes resulting in 5 measurements for each treatment (0min, 15min, 30min, 45min, 60min). Animal distribution data were tested for differences over time within each treatment. Respective ANOVAs that tested every treatment for differences between the subsequent measurements did not reveal any significant differences and data were therefore pooled for each treatment over time.

#### Swimming velocity

To determine adaptive swimming behavior, we conducted another experiment using ‘control’ and ‘fed *Utricularia*’ specimens. After preparing the treatments, the individuals were placed into a tank (12.5 cm × 10 cm × 2.5 cm) containing only charcoal-filtered tap-water (20°C ± 1°C) and were given five minutes for acclimation before the recordings began (27). We recorded the animals for five minutes at a frame rate of 30 fps using a Nikon D5100 (equipped with Nikon DX AF-S Nikkor 18-105 mm 1:3.5-5.6 G ED; Nikon Corporation, Tokio, Japan). Afterwards, we analyzed ca. 800 sequences of 5 seconds in which the animals were moving in a straight line and in parallel to the front pane of the tank. Movement of the animals’ geometric centers were tracked by hand using a self-scripted Matlab application (Matlab R2014b, The Mathworks Inc., Natick, MA, 2015). This program delivers the swimming velocity at any point in time and was subsequently used to calculate an average velocity for each individual. In total, we recorded and analyzed the swimming movements of 200 animals of each treatment.

#### Swimming mode

In *Daphnia* three different swimming modes can be classified: ‘hop & sink’, ‘zooming’ and ‘looping/spinning’ (47–51). The ‘hop and sink’ mode is characterized by alternating upward movements, powered by forceful strokes of the second antennae (hops), interrupted by periodical breaks (sink). In the ‘zooming’ mode daphniids display a series of fast swimming strokes with no sinking phases in-between. In comparison, the ‘hop and sink’ mode is a rather slow swimming mode (<10mm/sec) whereas the ‘zooming’ mode is rather fast (>15mm/sec) (48). The ‘looping/spinning’ mode is displayed as a series of backward loopings. From the aforementioned recorded videos, we randomly analyzed 65 videos per treatment and determined the proportions of the swimming modes ‘hop & sink’ and ‘zooming’ since these were the dominant movement patterns. That was done by randomly choosing a time frame of 30 seconds in each of these videos, in which the animal was clearly visible and swimming in parallel to the aquarium’s front pane. In all of the 65 videos per treatment a time frame meeting our requirements was found.

#### Predation experiments

We conducted predation trials to determine the effect of phenotypic changes on *U. australis* capture efficiency. For that, we placed 20 five-day-old ‘fed *Utricularia*’ or ‘control’ animals into a glass vial filled with 40 ml of charcoal-filtered tap water that contained a 5 cm long shoot of *U. australis* with a defined number of 30 empty traps. This setup was placed in a climate chamber at 20°C ±1°C and a day-night-cycle of 16:8 h. We counted the number of surviving animals twenty-four hours after the start of the experiment. We repeated the experiment 10 times for each treatment.

### Statistical Analysis

For the statistical analysis of our experimental data we used R x64 3.4.2(52) with a significance threshold ≤ 0.05 for all conducted tests. Tests and plots were conducted using the packages “ggplot2”(53), “gdata”(54), “ggpubr”(55), “ggsignif”(56), and “rstatix”(57).

Data of the 2D measurements followed a normal distribution (Shapiro test), so that we conducted a multivariate analysis of variance (MANOVA) with post-hoc test (Bonferroni-corrected pairwise t-test) to compare the four treatments across the six consecutive days of the experiment. We calculated η^2^ to estimate effect sizes based on the model used for the MANOVA.

Data of life-history parameters, i.e., brood sizes and the portion of sexually mature females did not follow a normal distribution and was thus analyzed using Kruskal-Wallis rank sum test (Bonferroni-corrected pairwise Wilcoxon-test) followed by determination of effect sizes using η^2^ for each day.

The 3D data were based on the computed comparisons of the averaged point positions, using a displacement vector approach and furthermore, the point translocations along the coordinate axes (refer to Horstmann *et al*. 2018 for details). We tested these axes-wise point translocations with Wilcoxon-tests at a significance level ≤ 0.01, conducted within the Matlab environment. Significance levels were adjusted for multiple testing based using False-Discovery-Rate (FDR) approach (58). This approach estimates the probability of declaring a not-differing feature as significantly different among all significant features, given as ‘q-value’. Finally, the 3D-forms of plant-exposed and control individuals were compared using confidence ellipsoids (40). We calculated the effect size Pearson’s *r* using R for each conducted Wilcoxon-test and averaged them (mean) for each analyzed axis.

The analysis of the predator avoidance experiment was based on the sections superimposed on the tank’s front pane. In order to analyze the vertical distribution of experimental animals we summed up the counts for each row of the grid. Likewise, we summarized the animal count of every column to analyze the horizontal distribution. Horizontal and vertical distributions were tested for differences between treatments independently using a Kruskal-Wallis rank sum test (Bonferroni-corrected pairwise Wilcoxon-test) for each time-point of the experiment. Since we did not find any differences between the time-points, we eventually averaged all observations of animal distributions per treatment over time. Additionally, we analyzed this data-set as a distribution offset from the tap water control. For this, every animal count in the volumes was given an identical ‘weight’, enabling the calculation of a ‘center of mass’ in the 2D distribution for every treatment (averaged over time) based on the vertical and horizontal distributions. Using a vector plot, we illustrate the offset of each treatment’s specific center of mass in reference to the control’s center of mass.

For the statistical analysis of swimming velocity and swimming mode, we conducted Kruskal-Wallis rank sum tests followed by Bonferroni corrected pairwise Wilcoxon-tests between the respective treatments. Finally, we calculated η^2^ to determining effect sizes.

## Results

### Trap entrance dimensions

*U. australis* trap entrance dimensions were determined as 495 µm (±166 µm SD) average height and 613 µm (±147 µm SD) average width (n=20 each). Therefore, the trap entrances are typically wider than high (ratio ∼1:1.23).

### Morphological alterations

#### 2D investigation

In our first experiment, we tracked two-dimensional morphometrics, i.e. body length and body width, of *C. dubia* over a period of 6 days in 4 different treatments (‘tap water control’, ‘*Ceratophyllum* control’, ‘unfed *Utricularia’*, ‘fed *Utricularia’*). We found a significant effect of time and treatment on *C. dubia*’s body lengths as well as a significant interaction (MANOVA; time: *F*=437.163, *DF*=6, *p*<0.001, η^2^=0.606; treatment: *F*=114.530, *DF*=3, *p*<0.001, η^2^=0.079; treatment × time: *F*=5.367, *DF*=17, *p*<0.001, η^2^=0.021) (Fig. 1, SI Appendix, Table S1, Table S2). As the animals grow over time, we focused our Bonferroni-corrected post hoc analysis on the differences between treatments within the individual days of the experiment and found the induced animals showing significant differences in comparison to the controls. We found no differences between the two control treatments (i.e. ‘tap water control’ and ‘*Ceratophyllum* control’). On day 1 of the experiment, we found the ‘unfed’ and ‘fed *Utricularia’* exposed treatments to be significantly different in body lengths (SI Appendix, Table S3). The ‘unfed *Utricularia’* treatment showed larger body length than the ‘fed *Utricularia’* treatment. This difference between the two *Utricularia*-exposure treatments was not observed on day 2, 3 and 4 but on day 5 and 6. From day 2 onwards the two control treatments showed significantly larger body length than the ‘fed *Utricularia’* treatment (SI Appendix, Table S3). From day 3 onwards the ‘tap water control’ showed significantly larger body lengths than the ‘unfed *Utricularia’* treatment. From day 5 onwards the body length of both *Utricularia*-exposure treatments were significantly smaller than the control treatments (SI Appendix, Table S3).

**Figure 1:**
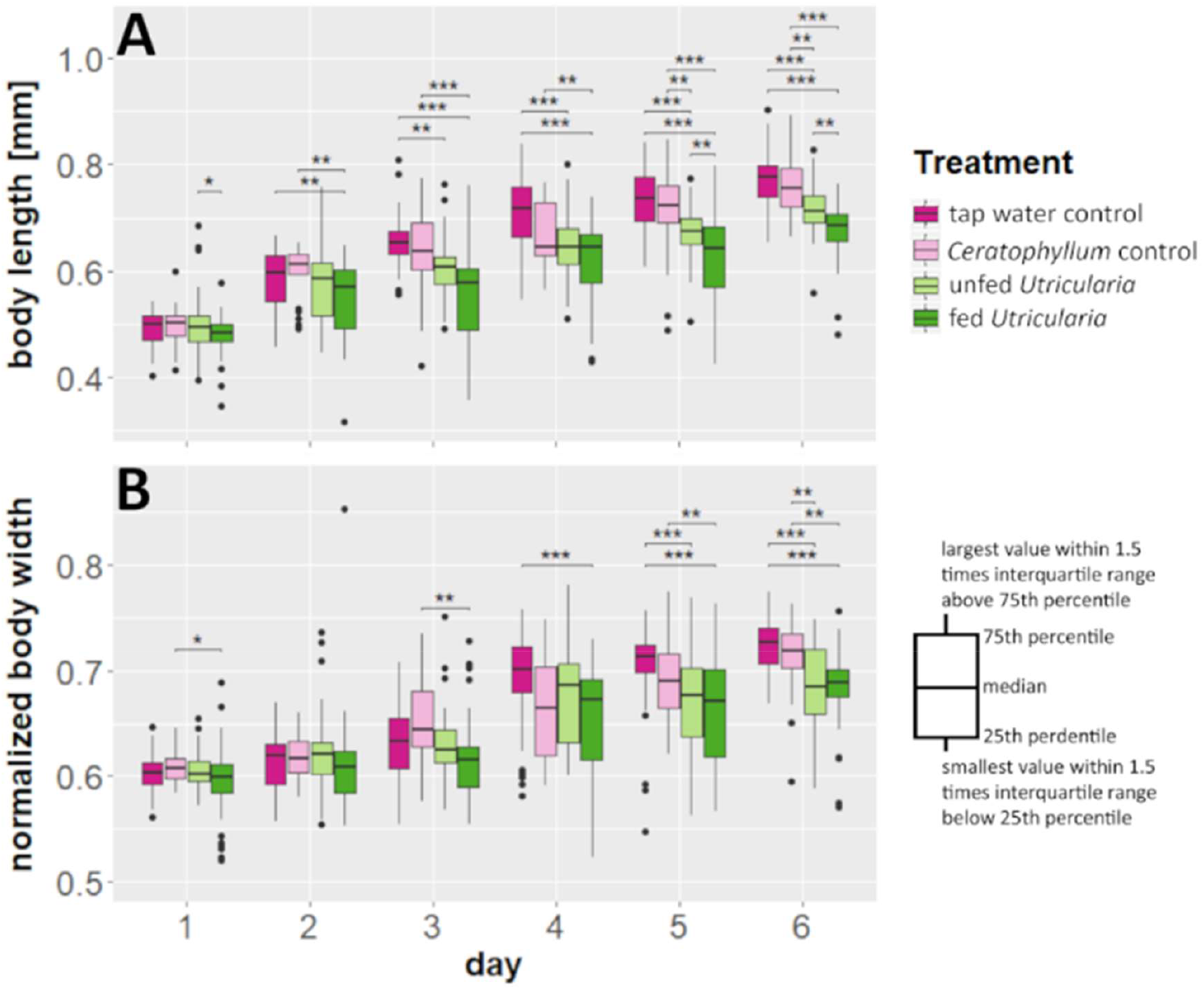
Morphological changes in *C. dubia* as a response to the presence of *U. australis*. (A) Body lengths measurements over a duration of 6 days for 4 different treatments including two control treatments and two Utricularia-exposed treatments. (B) Normalized body widths (body width/body length) accordingly. Utricularia-exposed animals show significantly smaller body length and normalized body width as the control treatments.

MANOVA of the normalized body width also revealed a significant effect of time and treatment as well as a significant interaction (time: *F*=253.224, *DF*=6, *p*<0.001, η^2^=0.508; treatment: *F*=41.110, *DF*=3, *p*<0.001, η^2^=0.041; treatment × time: *F*=4.789, *DF*=17, *p*<0.001, η^2^=0.027) (Fig. 1, SI Appendix, Table S1, Table S2). Using Bonferroni post hoc analysis, within the individual days but between the treatments, we neither detected significant differences between the two control treatments nor between the two *Utricularia*-exposure treatments. On day 1 and 3 of the experiment, a significant difference for normalized body widths was only detected between ‘fed *Utricularia*’ and the ‘*Ceratophyllum* control’ treatments with ‘fed *Utricularia*’ animals being smaller (SI Appendix, Table S4). For day 2, we found no significant differences between the treatments. On day 4, the normalized body widths of animals exposed to ‘fed *Utricularia*’ were significantly smaller than those of the ‘tap water control’ treatment (SI Appendix, Table S4). From day 5 onwards, morphological alterations became more prominent. On day 5 both *Utricularia*-exposed treatments had significantly smaller normalized body width than the ‘tap water control’ treatment (SI Appendix, Table S4). Furthermore, the normalized body width of the ‘fed *Utricularia*’ treatment was significantly smaller than that of the ‘*Ceratophyllum* control’ treatment on day 5 (SI Appendix, Table S4). On day 6 both *Utricularia*-exposed treatments showed significantly smaller normalized body width than the two control treatments (SI Appendix, Table S4).

#### 3D analysis

In order to obtain a comprehensive insight into the morphological adaptations of we analyzed 3D shape differences between control and *Utricularia*-induced 5 days old *C. dubia* using the method described by Horstmann et al.(40). Using this approach, we confirmed the same significant differences between control (Fig. 2A) and *Utricularia*-exposed animals in five-days old specimens (Fig. 2B). These differences in overall appearance (Fig. 2C) are supported by the confidence ellipsoid analysis, as it revealed no overlaps indicating the overall difference between both morphotypes (Fig. 2D). We found mean Pearson’s *r* effect sizes of 0.733 for the dorso-ventral body axis, 0.791 for the anterior-posterior body axis, and 0.556 for the lateral body axis.

**Figure 2:**
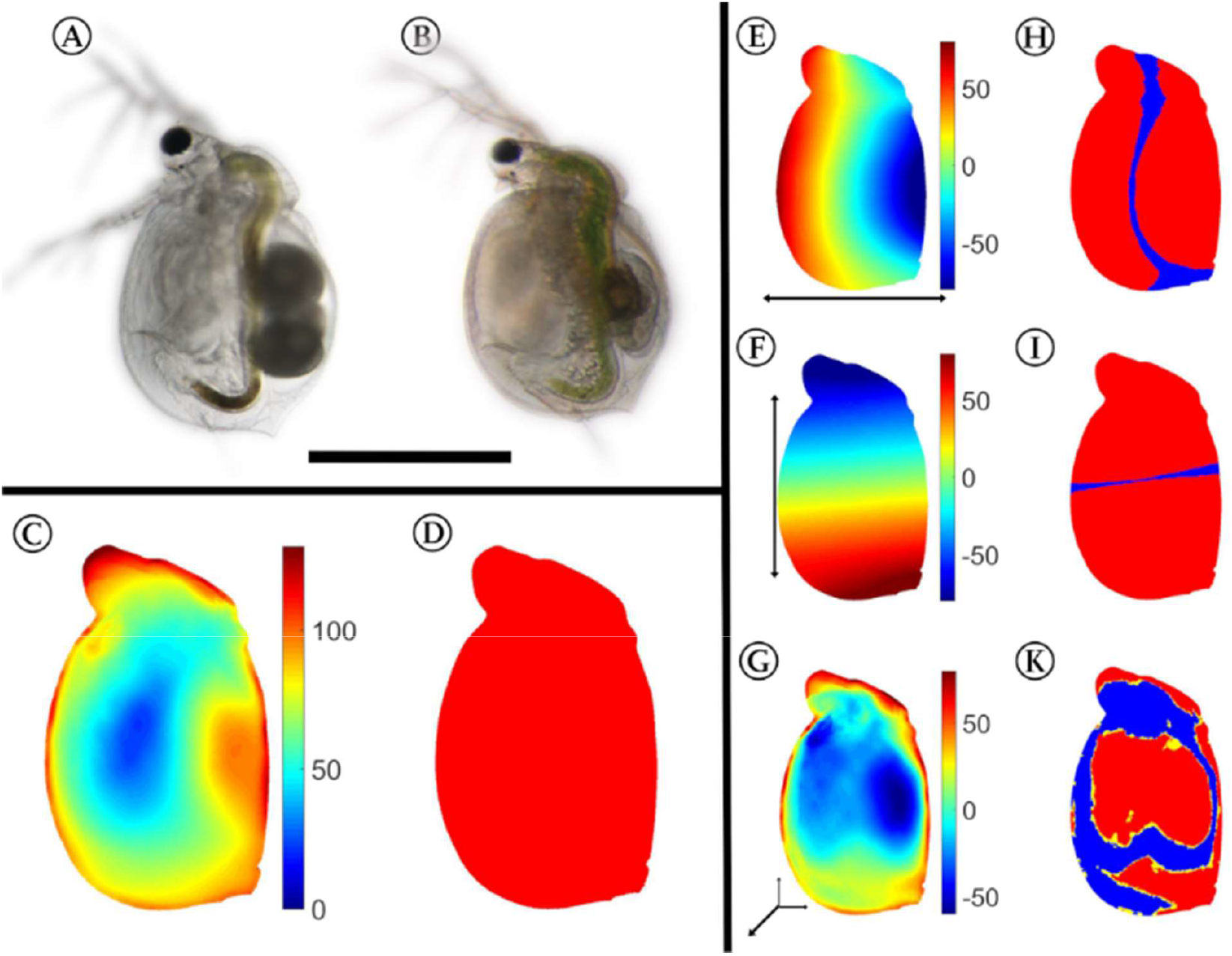
3D analysis of morphological alterations between control and Utricularia-exposed *C. dubia*. Control (A) and Utricularia-exposed *C. dubia* (B) of same age, scale bar = 1 mm. All subsequent analyses are projected on the average Utricularia-exposed animal. (C) Overall deformation, strong shifts are colored in shades of red, while small or no changes are indicated by shades of blue. (D) Confidence ellipsoid plot, revealing no overlapping confidence ellipsoids. (E)-(F): Here, shades of red indicate a shift in positive direction on that axis (dorsal/anterior/distal), shades of blue indicate a shift in negative direction on the respective axis (ventral/posterior/proximal). Shifts along the anterior-posterior (E) and dorso-ventral axis (F) show that the animals are smaller in the Utricularia-exposed morph. The deformation in the lateral dimension (G) gives regions of reduced and increased body width. Most of the found shifts are proofed significant with respective Wilcoxon tests and FDR-based q-values (H)-(K). These figures give regions with p-values of respective Wilcoxon tests lower than 0.01 coloured yellow, regions that showed in addition q-values lower than 0.001, are coloured red. For the respective analysis, all samples of both treatments were taken into account (ninduced= 8, ncontrol = 13).

*Utricularia-*exposed animals (Fig. 2B) are smaller than control animals (Fig. 2A) of the same age, which is manifested in the body length (ctrl=0.725 mm ±0.0175, induced=0.529 ± 0.038 mm, reduction of 27%). Additionally, we found a general dorsally directed shift of our landmarks of the ventral body regions (Fig. 2E). In contrast, our landmarks of the dorsal region are shifted ventrally, which makes the *Utricularia*-exposed animals slimmer than the controls. While the form alterations of the ventral and dorsal body regions are validated significant based on Wilcoxon-tests and FDR-testing, most shifts of our landmarks of the tail-spine region were not supported statistically significant (*p*>0.01, *q*>0.001) by the FDR approach (Fig. 2H).

Considering the shifts along the anterior-posterior body axis (Fig. 2F), we see that our landmarks of the anterior body parts (e.g. head) are significantly (*p*<0.01, *q*<0.001) shifted in posterior direction, while the posterior ones are significantly (*p* <0.01, *q*< 0.001) shifted in anterior direction (Fig. 2I).

In lateral direction, the strongest altered regions are the head, neck and brood pouch (Fig. 2G). Interestingly, the head’s lateral width is larger by about 90 µm (37%), leading to a total lateral width of 365 µm. The neck region’s width is larger by about 120 µm (35%), leading to a total width of 475 µm. This is mostly due to very pronounced fornices that are visibly formed only in *Utricularia*-exposed animals. In the region of the brood pouch, the *Utricularia*-exposed animals are thinner by about 90-120 µm. In the region of the second antenna joint, the *Utricularia*-exposed animals are laterally slimmer as well.

### Life-history shifts

We additionally determined the number of egg-carrying females as well as the individual clutch sizes of the animals used in our first experiment. Kruskal-Wallis tests revealed no significant differences in number of egg carrying females between the treatments (Fig. 3, SI Appendix, Table S5). However, from day 4 onward the ‘tap water control’ treatment showed a significantly larger clutch size than both *Utricularia*-exposed treatments (Day 4: χ^2^=317.54, df=3, p-value<0.001, η^2^=0.157; Day 5: χ^2^=586.6, df=3, p-value<0.001, η^2^=0.119). On day 6 the ‘*Ceratophyllum* control’ treatment also deposited significantly more eggs than the *Utricularia*-exposed treatments (Fig. 3, SI Appendix, Table S5) (Day 6: χ^2^=418.64, df=3, p-value<0.001, η^2^=0.137).

**Figure 3:**
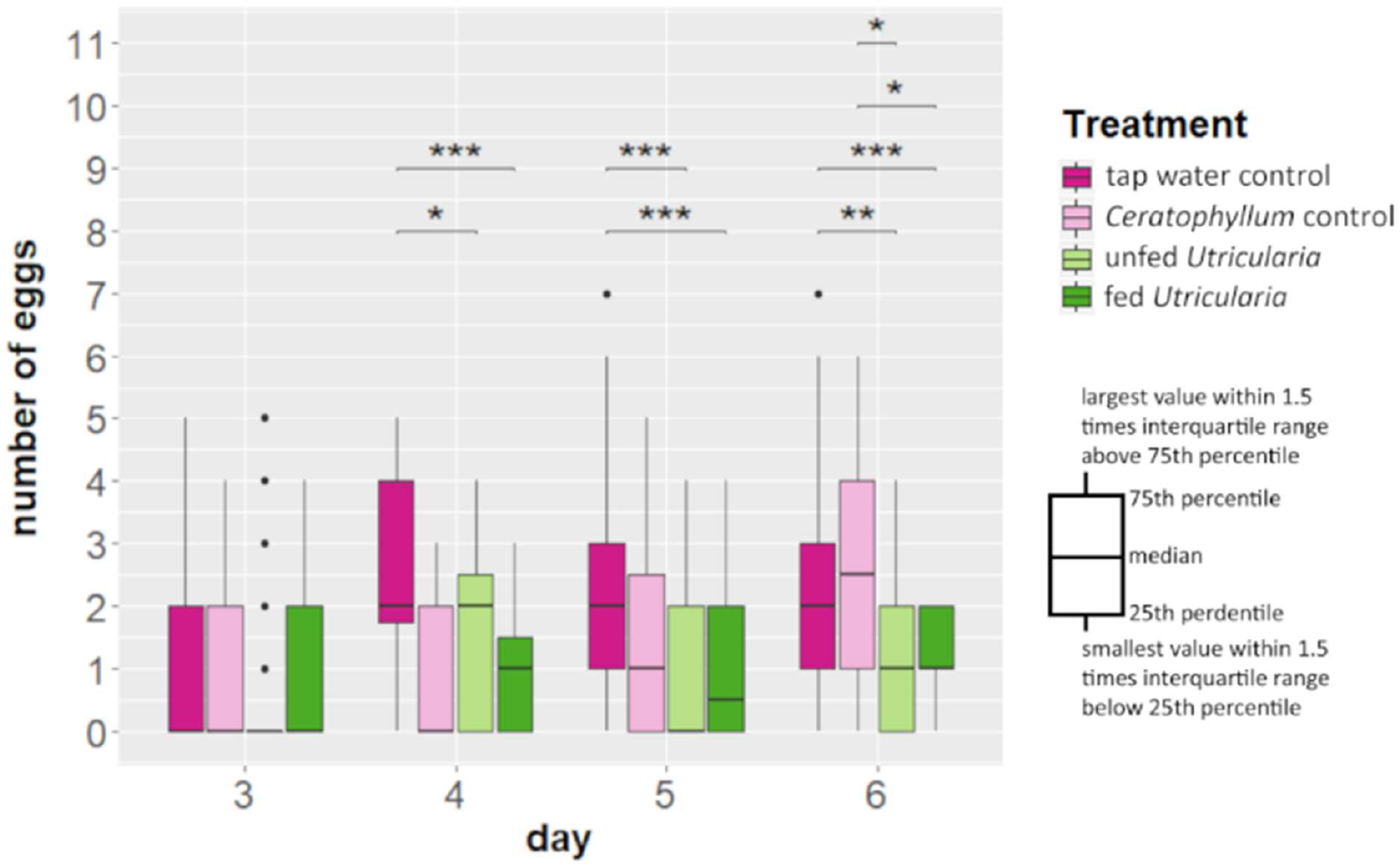
Changes in life history (in terms of clutch size alterations) of *C. dubia* in the presence of *U. australis*. *C. dubia* revealed smaller clutch sizes (p≤0.01) in presence of *U. australis* compared to the control treatments from day 4 onwards, stagnating at about 1 egg per female.

### Behavioral alterations

#### Predator avoidance

As a predator avoidance experiment we introduced 10 *C. dubia* into one of 5 different treatments (control animals/no plant, control animals/*Elodea*, control animals/*Utricularia*, ‘fed *Utricularia’* induced animals/*Elodea* and ‘fed *Utricularia’* induced animals/*Utricularia* and recorded their position using a grid of 6 by 3 squares drawn on front panes of the tanks. During the experiment, the majority of animals were observed to aggregate in the two upper edges of the tank (Fig. 4; ANOVA, F=5.265, p<0.001, η^2^=0.016), i.e. the top sections (Fig. 4B) of the most left and right columns (Fig. 4A). This was true for all treatments. In the control treatments (tap water controls as well as control animals facing *E. canadensis*) no side preference was observed. In all treatments including *U. australis* (*Utricularia*-exposed animals as well as control animals) a significant side preference away from the plant and towards the water surface was observed (Fig 4; ANOVA, F=10.260, p<0.001. η^2^=0.036).

**Figure 4:**
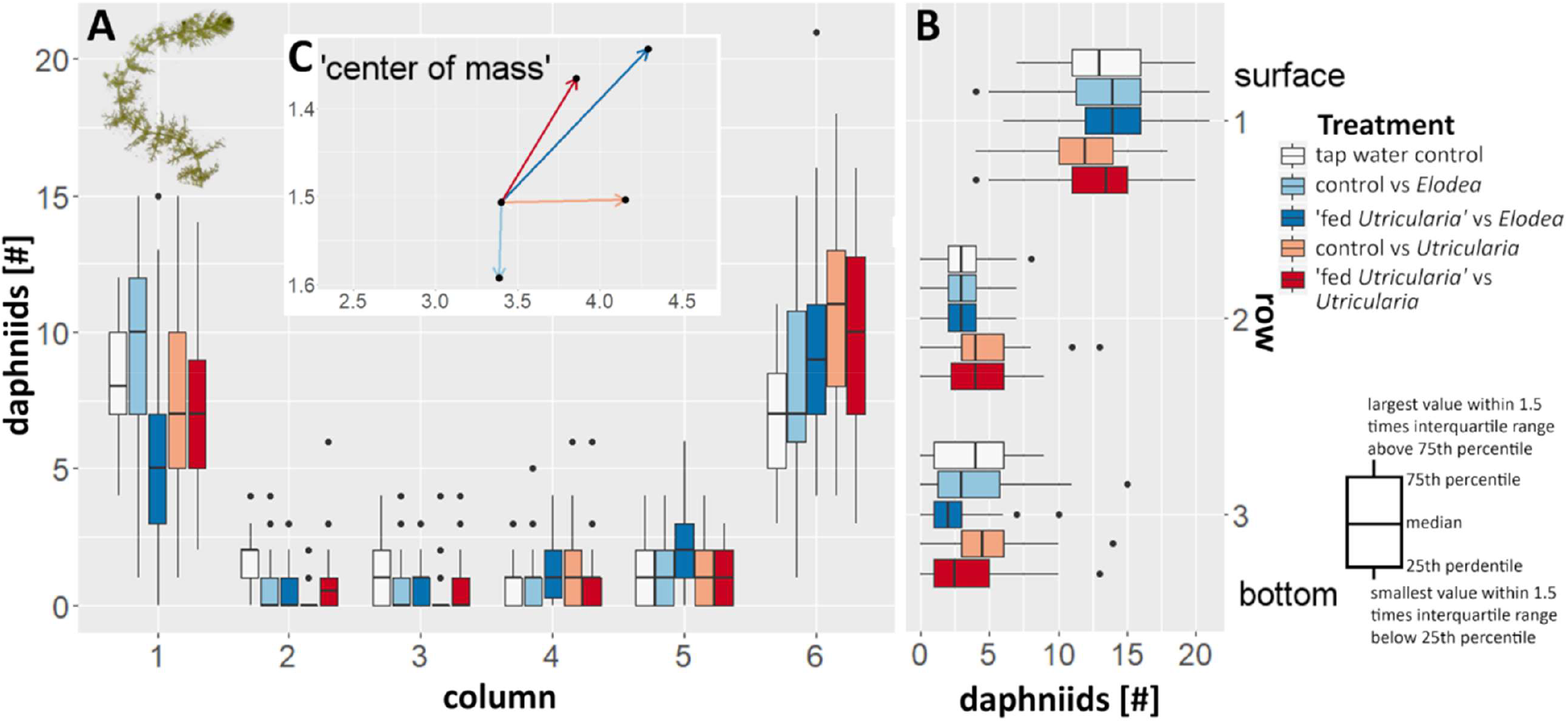
Changes in behaviour in *C. dubia* observed as averaged distribution pattern with respect to the presence of either *U. australis* or *E. canadensis*. (A) The box plots indicate the number of animals per column in the canvas drawn on the tank front pane. Increasing numbers on the x axis are equal to an increase in distance to the respective plant used in that treatment (1 equals to the same column as the plant, 6 is the opposite tank side). (B) The box plots indicate the number of animals per row. (C) The additional vector graph inlet is indicating the average positioning of the animals in respective to the plant by depicting a vector that represents the ‘calculated center of mass’ for every treatment as an offset from the tap water control-treatment.

#### Swimming modes

In order to detect differences in swimming behavior, we recorded videos of swimming *C. dubia* in control and ‘fed *Utricularia*’ states. We analyzed the recorded videos for the proportion of swimming modes that occur in the control and induced treatment. We found significant differences between the treatments (Kruskal-Wallis rank sum test; *chi-squared*=53.978, *DF*=3, *p*≤0.001, η^2^_Hop and Sink_=0.028, η^2^_Zooming_ =0.023). In the tap water control there was no significant difference between the percentage of duration of ‘zooming’ and ‘hop & sink’ swimming mode (Bonferroni corrected pairwise-Wilcoxon-test; *p*≤0.05). The tap water control animals performed ‘zooming’ in 43% of the investigated time, whereas ‘hop & sink’ was performed 57% of the time. The animals of the ‘fed *Utricularia*’ treatment showed significant differences in percentage of duration of the used swimming mode (Bonferroni corrected pairwise-Wilcoxon-test; *p*≤0.001). They performed ‘zooming’ in roughly 25% of the time and ‘hop & sink’ in about 75% of the time (Fig. 5A). This resulted in significant higher percentage of ‘hop & sink’ mode in the ‘fed *Utricularia*’ treatment in comparison to the tap water control and significant lower respective percentage in ‘zooming’ mode (Bonferroni corrected pairwise-Wilcoxon-test; *p≤*0.05 for both comparisons).

**Figure 5:**
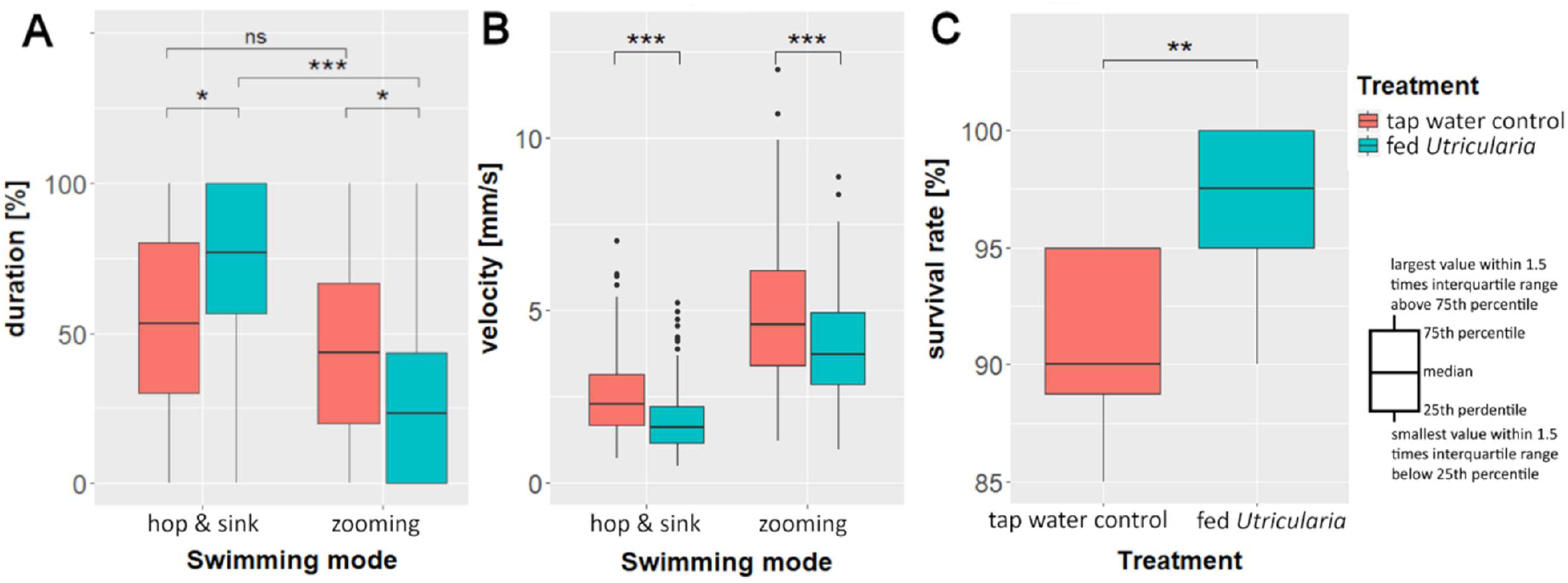
Behavioral changes in *C. dubia* as response to the presence of *U. australis*: average duration (A) and velocity (B) of the two observed swimming modes in the ‘swimming modes’ experiments. (C) Survival rate of 20 five-days old *C. dubia* (either control or Utricularia-exposed) over 24h in the presence of 30 *U. australis* traps.

#### Swimming velocity

We additionally analyzed the above-mentioned video recordings for *C. dubia’s* swimming speed. Our analysis of the average swimming velocity showed that induced animals swam significantly slower than the control animals (Kruskal-Wallis rank sum test; *chi-squared*=359.09, *DF*=3, *p*≤0.001, η^2^_Hop and Sink_=0.06, η^2^_Zooming_ =0.025). That is true for both swimming modes (pairwise Wilcoxon test; *p*_hop& sink_≤0.001; *p*_zooming_≤0.001). In the ‘hop & sink’ mode, the average (median) velocity of control animals was ∼2.3 mm*s^-1^ whereas that of induced animals was 1.6 mm*s^-1^. In the ‘zooming’ mode, the control animals swam at velocities of 4.6 mm*s^-1^, while the induced animals swam with an average (median) velocity of 3.7 mm*s^-1^ (Fig. 5B). Resulting in a much shorter swimming distance over time for induced animals in either swimming mode.

#### Predation experiments

Finally, we performed a predation experiment in order to compare survival chances of control and ‘fed *Utricularia’* individuals of *C. dubia*. The analysis of the predation data revealed significant differences between the two treatments (Mann-Whitney U-test; *U*=10.5, *p* ≤ 0.05, *n*=10, *r*=0.691). More induced animals (median survival rate 97.5%) than control animals (median survival rate 90%) survived in the presence of *U. australis*. The maximum number of trapped animals in our experiments were four for undefended and two for defended animals (Fig. 5C). In other words, *U. australis* caught every 10^th^ control animal but only every 40^th^ induced animal.

## Discussion

In this study, we observed predator-induced, phenotypically plastic responses in form of morphological, life history and behavioral traits of *C. dubia* exposed to *U. australis*. The two species, both representatives of cosmopolitan clades, are members of a naturally co-occurring predator-prey system native to Central Europe (29–31).

Phenotypic plasticity in plant-animal interactions is long known and especially herbivore-induced plant defenses are well studied (59). Furthermore, herbivores are described to express dietary-induced plasticity in morphology and behavior allowing to deal with plant defenses (60,61). However, to our knowledge plant-induced defenses in an animal have not been described yet. In the following, we discuss the observed plastic responses and their adaptive benefit together with first insights into the nature and origin of the eliciting cue(s).

### Morphological adaptations

We observed a change in overall body shape in *C. dubia* when exposed to *U. australis*: the animals are shorter and slimmer in the dorso-ventral dimension but functionally increase their lateral size via elongation of their fornices compared to control animals of the same age. The latter is substantial as their lateral dimension increases by 37%. Given the apparent gape limitation of the bladderwort suction traps we hypothesize that the defensive mechanism is a combination of functional size increase and suction force reduction at the same time. The elongated fornices can hinder the animal’s entry into the trap by interfering with the traps entrance gape size while the slim body simultaneously allows the surrounding water to freely flow into the trap and eventually the pressure difference to equalize. We assume that the latter is key to this defense strategy since an overall increase in body dimensions would lead to a total or nearly total blockage of the trap entry with the result that the animal’s body would experience the (nearly) full amount of lethal suction forces produced by the trap. Based on our data, induced animals will only be able to block the smaller trap entrances (lateral dimension: 475 µm; smallest trap entrances: 495±166 µm). As inducible morphological defenses in daphniid species are known to continuously grow with every molting cycle, we are certain that our data merely represent the threshold of the defensive effect and with continuous molts the defensive effect will increase. Additional to the aforementioned blocking effect the slimmer body may reduce the chances to mechanically trigger the traps. Smaller animals may also face smaller drag forces which could increase survival chances by reducing the acceleration of the animal towards the trap once the trap is triggered. Any of these effects may also explain the prey preference towards larger prey as reported for two other *Utricularia* species by Guiral & Rougier (62).

### Life-history adaptations

In animals, exposed to *U. australis* during development, we detected changes of life-history parameters as they produce significantly less offspring per brood. Such a reduced number of offspring has been reported as a defense against visually hunting fish (63). *D. magna* becomes more prone to visual predator detection the more eggs are deposited in the brood pouch (64). In case of the mechanosensory-dependent *U. australis* predation, it is unlikely that a reduced number of offspring has the aforementioned effects. It is more likely to represent costs associated with the expression of defenses and the material required for the elongated fornices, and/or of a smaller brood pouch caused by the shape alteration. The decreased somatic growth rate may limit the amount of food that can be ingested since a reduced body size also limits the food filtration. Therefore, the observations may also be explained by the size-efficiency hypothesis (65).

### Behavioral adaptations

In comparison to morphological and life history adaptations that require some time to be expressed (here up to 5 days) (10,15,51,66,67), behavioral responses can be expressed quickly (6,27). Behavioral defenses, especially in *Daphnia*, can therefore function as temporary defenses, that bridge the time lag between predator perception and morphological defenses expression (68). In the presence of the carnivorous plant, the behavioral and morphological changes of *C. dubia* are expressed simultaneously. Maybe, the morphological changes alone do not suffice against a very effective predator like *U. australis*, which can have a capture rate of 100% for undefended *C. dubia* in different juvenile instars (32).

In our experiments, *Utricularia*-exposed *C. dubia* avoided the presence of *U. australis* and *C. demersum*, likewise. Animals of the control group avoided only *U. australis*. Potentially, *Utricularia*-exposed animals show higher alertness that makes them avoid any regions shaded by plants. This might be an alteration in phototactic behavior as only our treatments that directly faced *U. australis* or were exposed to it prior to the experiment showed significant ‘open water’ preferences. Control animals showed no significant avoidance of shaded areas. Fish evoke similar but opposite behavioral responses in *D. magna*: Lauridsen & Lodge (69) demonstrated that *D. magna* seeks shelter in plant thickets when threatened by young sunfish (*Lepomis cyanellus*).

In our analysis of swimming modes and speed we found the ‘hop & sink’ mode, which is a less directed, slower movement significantly increased in induced animals. Additionally, we found significantly reduced velocities of both observed modes in the *Utricularia*-exposed treatments compared to the control treatments. This overall reduction in swimming speed will either reduce the encounter rate between predator and prey (70) and/or it reduces the possibility to activate the trigger hairs on the *U. australis* trap door by reducing the kinetic energy of the animals (36). Such a behavioral adaptation is also known from *D. magna*, which reduce their swimming velocity in the presence of fish cues or homogenized conspecifics (27,71). A reduced swimming speed often comes at the cost of reduced feeding, which eventually leads to a reduced growth and fecundity (70).

### Predation trials

In our predation trials we tested if the above-described defenses are beneficial and render *C. dubia* less susceptible to this plant predator. We show that induced animals expressing behavioral and morphological defenses are less often captured and thus are better protected against *U. australis*. As these phenotypic changes increase the survival of *C. dubia*, we hypothesize that they evolved in response to *U. australis* predation. The increase of survival rate of 7.5% in induced animals may seem rather insignificant on first sight, but it means for *Utricularia* to catch only every 40^th^ daphniid instead of every 10^th^. Additionally, it is safe to assume that we only tested the early defensive effect in ontogeny as these defensive structures grow even more pronounced over subsequent molts like it is described for several daphniid species (e.g.(15)). However, this remains to be investigated in future studies. The defensive effect appears to be based on a synergism of behavioral and morphological adaptations with simultaneous life-history changes reflecting trade-offs of both types of defenses.

### Origin of the defense-inducing stimulus

The origin of the cue that induces the observed alterations in *C. dubia’s* morphology, life-history and behavior is unclear. Based on our experiments we cannot exclude that *C. dubia* could sense *U. australis* trap firings via mechano-receptors or identify the plant optically. However, we suggest that *C. dubia* detects chemical substances released by *U. australis*. Since *U. australis* and *C. dubia* were separated by net cages in our experiments, mechanical and visual cues were strongly damped while chemical cues were not. Also, there are many examples described, especially in *Daphnia*, where predator presence is detected chemically (44,72). Furthermore, we found reactions of *C. dubia* not only in fed *U. australis* treatments but also in unfed *U. australis* treatments. This suggests that it is not an alarm cue from conspecifics but a chemically active substance, a kairomone (73), released by *Utricularia* but not directly connected to predation activity. In contrast to this, the kairomone of *Chaoborus* larvae is released with digestive liquids (8,74) and induces neckteeth formation in *D. pulex* (75) only if predators are feeding. Nonetheless, *U. australis* fed with conspecifics of the investigated *C. dubia* induced stronger responses, e.g., stronger reduction of body length (Fig. 1). This suggests that the cue is stronger with successful capture or, at least, higher trap activity. For arming the traps, *U. australis* bladders constantly pump water out of their interiors (33) (for which the mechanism and pathway are not yet fully understood). They also exhibit spontaneous firings once a critical negative pressure is achieved (76). Moreover, prey capture leads to increased plant growth and production of larger traps (77) with a higher spontaneous firing rate (and thus resetting rate) (78). If *C. dubia* is able to sense (spontaneous) trap firings, detect the processes of digestion (79), trap resetting, respiration rate (80) and/or water excretion, *C. dubia* would have indirect measure(s) not only of trap presence but also activity. From *Daphnia*, it is known that they react to predator kairomones but also to broadly defined alarm signals (81). Alarm cues appear unlikely in our case given that prey organisms are not wounded during ingestion. In order to clarify the origin of the cue, further experiments are needed.

## Conclusion

Predator induced phenotypic plasticity is discussed to evolve under certain circumstances (4): First, the predation pressure must be variable and occasionally strong. Second, the predator must be perceptible by a reliable cue. Third, the induced defense must be effective. Fourth, the defense should be associated with costs or trade-offs. *U. australis* shows variability in abundance throughout the year with high abundances during summer and a resting stage during winter (82). Furthermore, the trap number of *U. australis* varies in dependence of biotic and abiotic factors reaching peak densities that pose severe threat to zooplankters (83). Given that *U. australis* exhibits the necessary variability in trap abundance and causes high predatory pressure at least during the summer months, the first prerequisite for inducible defenses is already fulfilled. Second, we present strong evidence for a reasonably reliable cue that enables *C. dubia* to perceive *U. australis* and react on its presence with a set of behavioral and morphological alterations. Third, we show that these adaptive changes are effective as induced *C. dubia* are consumed less by *U. australis*. Fourth, our experiments also show that the fecundity of induced animals is reduced, thus these alterations come at the expense of population growth rate. In summary, our study strongly suggests the evolution of animal inducible defenses against a predatory plant.

With inducible defense strategies being highly predator-specific and the fact that *U. australis* is only one representative of a cosmopolitan genus containing more than 250 species, we expect *C. dubia* not to be the only member of the Daphniidae family to thwart this “green threat” with inducible defenses. The carnivorous waterwheel plant (*Aldrovanda vesiculosa*) with snap-traps is another aquatic predator for daphniids and other zooplankters (84,85). In fact, given the variety of carnivorous plants, their trapping principles and sometimes narrow prey spectra (86) there probably is a number of inducible defenses against them yet to be identified in different species and ecosystems.

The predation experiment data did not follow a normal distribution and was therefore analyzed using non-parametric methods. The treatments were tested for differences using a Mann-Whitney U-test followed by a calculation of Pearson’s *r* for effect size.

## Author Contributions

SK, MH, SP, TS and RT designed the experiments. EB and NK conducted the experiments with supervision of SK and MH. LCW and RT supervised the culturing of *C. dubia*, SP and TS supervised the culturing of *U. australis*. SK and MH analyzed the data. LCW, SP and AW revised the statistical analysis. SK, MH and LCW wrote the first draft, SP, AW, TS and RT revised the draft and equally contributed to the final draft. TS and RT supervised the study and contributed their expertise and laboratory facilities.

