## Supplementary material for "Facing the green threat: A waterflea’s defenses against a carnivorous plant": SI Appendix

**Supplementary Materials**

**Table S1: Number of total animals measured in the 10 replicates of the 6-day experiment as well as the ANOVA F-values of the respective statistics (Fig 1).**

| **Day** | **1** | | | **2** | | | **3** | | | **4** | | | **5** | | | **6** | | |
|---|---|---|---|---|---|---|---|---|---|---|---|---|---|---|---|---|---|---|
| | **n** | **ANOVA F-value** | | **n** | **ANOVA F-value** | | **n** | **ANOVA F-value** | | **n** | **ANOVA F-value** | | **n** | **ANOVA F-value** | | **n** | **ANOVA F-value** | |
| **Treatment** | | Body length | Body width | | Body length | Body width | | Body length | Body width | | Body length | Body width | | Body length | Body width | | Body length | Body width |
| tap water control | 84 | 3.38 | 3.07 | 69 | 6.6 | 1.14 | 45 | 14.15 | 4.52 | 63 | 17.42 | 6.45 | 73 | 32.95 | 14.72 | 55 | 29.21 | 17.25 |
| *Ceratophyllum* | 37 | | | 30 | | | 35 | | | 25 | | | 44 | | | 29 | | |
| fed *Utricularia* | 79 | | | 72 | | | 42 | | | 50 | | | 50 | | | 43 | | |
| unfed *Utricularia* | 69 | | | 59 | | | 38 | | | 49 | | | 64 | | | 43 | | |

**Table S2: Mean body length and body width in mm ±SD of *C. dubia* over the 6 days of the experiment.**

| Treatment | Day 1 | | Day 2 | | Day 3 | | Day 4 | | Day 5 | | Day 6 | |
|---|---|---|---|---|---|---|---|---|---|---|---|---|
| | Body length | Body width | Body length | Body width | Body length | Body width | Body length | Body width | Body length | Body width | Body length | Body width |
| **tap water control** | 0.49±0.03 | 0.30±0.02 | 0.59±0.05 | 0.36±0.04 | 0.66±0.05 | 0.41±0.04 | 0.71±0.07 | 0.49±0.07 | 0.74±0.06 | 0.52±0.05 | 0.77±0.05 | 0.56±0.05 |
| ***Ceratophyllum*** | 0.50±0.03 | 0.30±0.02 | 0.60±0.05 | 0.37±0.04 | 0.64±0.07 | 0.42±0.07 | 0.67±0.06 | 0.45±0.07 | 0.71±0.07 | 0.49±0.07 | 0.76±0.06 | 0.54±0.06 |
| **fed *Utricularia*** | 0.48±0.03 | 0.29±0.02 | 0.55±0.07 | 0.33±0.05 | 0.57±0.09 | 0.35±0.08 | 0.62±0.07 | 0.41±0.07 | 0.63±0.08 | 0.42±0.08 | 0.67±0.06 | 0.46±0.06 |
| **unfed *Utricularia*** | 0.50±0.05 | 0.30±0.03 | 0.57±0.07 | 0.36±0.05 | 0.60±0.06 | 0.38±0.04 | 0.65±0.06 | 0.44±0.06 | 0.67±0.05 | 0.45±0.05 | 0.71±0.05 | 0.49±0.05 |

**Table S3: Bonferroni corrected pairwise t-test results for 2D morphology body length analysis. Significant results marked in red.**

| Day | 1 | | | 2 | | | 3 | | |
|---|---|---|---|---|---|---|---|---|---|
| | Tap water ctrl | fed *Utric.* | *Cer.* ctrl | Tap water ctrl | fed *Utric.* | *Cer.* ctrl | Tap water ctrl | fed *Utric.* | *Cer.* ctrl |
| fed *Utricularia* | 0.197 | | | 0.0022 | | | 4.9e-8 | | |
| *Ceratophyllum ctrl* | 1 | 0.176 | | 1 | 0.0013 | | 1 | 4.5e-5 | |
| unfed *Utricularia* | 1 | 0.023 | 1 | 1 | 0.187 | 0.3159 | 0.0042 | 0.0954 | 0.1835 |
| Day | 4 | | | 5 | | | 6 | | |
| | Tap water ctrl | fed *Utric.* | *Cer.* ctrl | Tap water ctrl | fed *Utric.* | *Cer.* ctrl | Tap water ctrl | fed *Utric.* | *Cer.* ctrl |
| fed *Utricularia* | 2.8e-10 | | | <2e-16 | | | 2.6e-14 | | |
| *Ceratophyllum ctrl* | 0.1649 | 0.0077 | | 0.2034 | 6.8e-9 | | 1 | 4.2e-9 | |
| unfed *Utricularia* | 5.3e-5 | 0.1474 | 0.9692 | 2.7e-8 | 0.0025 | 0.0079 | 5e-6 | 0.0066 | 0.0029 |

**Table S4: Bonferroni corrected pairwise t-test results for 2D morphology body width analysis. Significant results marked in red.**

| Day | 1 | | | 2 | | | 3 | | |
|---|---|---|---|---|---|---|---|---|---|
| | Tap water ctrl | fed *Utric.* | *Cer.* ctrl | Tap water ctrl | fed *Utric.* | *Cer.* ctrl | Tap water ctrl | fed *Utric.* | *Cer.* ctrl |
| fed *Utricularia* | 0.34 | | | 1 | | | 1 | | |
| *Ceratophyllum ctrl* | 1 | 0.04 | | 1 | 1 | | 0.056 | 0.0026 | |
| unfed *Utricularia* | 1 | 0.16 | 1 | 1 | 0.52 | 1 | 1 | 0.7674 | 0.2504 |
| Day | 4 | | | 5 | | | 6 | | |
| | Tap water ctrl | fed *Utric.* | *Cer.* ctrl | Tap water ctrl | fed *Utric.* | *Cer.* ctrl | Tap water ctrl | fed *Utric.* | *Cer.* ctrl |
| fed *Utricularia* | 0.00024 | | | 7.9e-8 | | | 5.2e-8 | | |
| *Ceratophyllum ctrl* | 0.06692 | 1 | | 0.1841 | 0.0085 | | 1 | 0.001 | |
| unfed *Utricularia* | 0.62437 | 0.09849 | 1 | 3.3e-6 | 1 | 0.1059 | 4.8e-7 | 1 | 0.004 |

**Table S5: Mean number off eggs ±SD for each treatment on day 3, 4, 5 and 6.**

| Treatment | Day 3 | Day 4 | Day 5 | Day 6 |
|---|---|---|---|---|
| **tap water control** | 0±1.66 | 2±1.51 | 2±1.65 | 2±1.86 |
| ***Ceratophyllum*** | 0±1.20 | 0±1.18 | 1±1.71 | 2.5±1.73 |
| **fed *Utricularia*** | 0±1.11 | 1±1.00 | 0.5±1.14 | 1±0.68 |
| **unfed *Utricularia*** | 0±1.71 | 2±1.45 | 0±1.20 | 1±1.13 |
